# The difference in immunohistochemical reactivity of monoclonal antibodies against amino-terminal residues of amyloid-β peptide

**DOI:** 10.1101/2025.10.15.682678

**Authors:** Kana Araki, Kenta Yamauchi, Shogo Ito, Masato Koike, Hioki Hiroyuki

## Abstract

Immunohistochemistry for amyloid-β (Aβ) peptide is an indispensable method for Alzheimer’s disease (AD) research. Despite a wide variety of available antibodies against the peptides, the difference of immunohistochemical reactivity is not fully described among anti-Aβ antibodies. Immunohistochemical reactivity of Abs against Aβ peptides is critical for accurate and reliable evaluation of Aβ burden in patients as well as models of AD. Here, we examined immunohistochemical reactivity of two mouse and one rabbit monoclonal antibodies against Aβ N-terminal regions using two AD mouse models, *App*^*NL-F*^ and *App*^*NL-G-F*^. 6E10, 82E1 and D54D2 Aβ antibodies were used in this study. We found significant differences in the immunohistochemical reactivity in both *App*^*NL-F*^ and *App*^*NL-G-F*^ models. While 6E10 immunoreactivity was mainly localized to Aβ plaques, D54D2 and 82E1 antibodies stained much more broadly beyond plaques. Interestingly, the latter two Abs showed blurred filamentous immunoreactivity beyond amyloid plaque cores. Double immunostaining using a tyramide signal amplification method, Fluorochromized Tyramide-Glucose Oxidase (FT-GO), suggested that the differential immunohistochemical outcomes were only partially attributable to their sensitivity. Moreover, heat induced epitope retrieval (HIER) did not affect the differential immunohistochemical outcomes. Our analysis indicates that outcomes of Aβ immunohistochemistry highly depends on the antibody used in the study.

## Introduction

A major histopathological hallmark of Alzheimer’s disease (AD) is deposition of amyloid-β (Aβ) peptides in the brain (1). Aβ peptide is produced through proteolytic processing of a transmembrane protein, amyloid precursor protein (APP) (2, 3). While β-secretase cuts on the N-terminal end of the Aβ peptide, γ-secretase cleaves APP protein at multiple sites at the C-terminal ends to give rise to heterogeneous Aβ species. Aβ peptide is highly fibrillogenic and aggregates into higher-order species such as oligomers, protofibrils, fibrils and plaques, which have detrimental effects on brain integrity (2, 3). More than 30 mutations in APP gene have been reported to cause familial Alzheimer’s disease (FAD) (4). Aβ pathology precedes the clinical onset of dementia by over 20 years (5). These findings support the amyloid cascade hypothesis positing that deposition of Aβ plaques in the brain triggers a cascade of events that leads to neurodegeneration (6, 7). The recent successes of Aβ-immunotherapy trials further strengthen the hypothesis (8).

Among multiple Aβ imaging techniques, immunohistochemistry (IHC) with antibodies (Abs) against Aβ peptides is superior in sensitivity and specificity (9) and widely used for measuring Aβ load in AD patients as well as models. Microscopic examination of IHC staining for Aβ peptides is further utilized for the definitive diagnosis and neuropathological staging of AD in postmortem brains (9, 10). Various kinds of Abs against Aβ peptides have been developed over the past decades and adapted for IHC analysis (11-16). They show differences in clonality, host species, target amino acid sequences and modification. However, there is no consensus on which Abs should be used for the detection of Aβ peptide by IHC. Critically, the difference in IHC reactivity among anti-Aβ Abs are not fully described. The differential IHC reactivity of the Abs should affect neuropathologic analysis of Aβ deposition by IHC staining.

Here, we examined IHC reactivity of two mouse monoclonal and one rabbit monoclonal Abs against N-terminal residues of human Aβ peptides using two *App* knock-in (KI) mouse models. These Abs showed significant differences in both mouse models regarding IHC outcomes. We further asked whether the differences could be attributable to their sensitivity and antigen masking.

## Materials and Methods

### Animals

This study was carried out in strict accordance with Fundamental Guidelines for Proper Conduct of Animal Experiments by the Science Council of Japan (2006). All animal procedures were conducted in compliance with ARRIVE (Animal Research: Reporting *In Vivo* Experiments) guidelines. The protocol was approved by the Institutional Animal Care and Use Committees of Juntendo University (Approval No. 2023194). All surgery was performed under sodium pentobarbital anesthesia, and all efforts were made to minimize suffering.

Male *App*^*NL-G-F/NL-G-F*^ (17) mice (RBRC06344, RIKEN BioResource Research Center) at 14 months old were used. The mice were housed in 12/12 h light/dark cycle at 20–25 °C under specific pathogen-free conditions and given access to food and water ad libitum. Mice were euthanized if they reached the humane endpoints: self-mutilation, breathing problems/cyanosis or inability to access food or water. Animal health and behavior were monitored twice a week. Three randomly selected mice were subjected to immunohistochemical examination.

### Tissue preparation

Anesthesia was induced by intraperitoneal injection of an overdose of sodium pentobarbital (200 mg/kg; P0776, Tokyo Chemical Industry) in accordance with American Veterinary Medical Association (AVMA) Guidelines for the Euthanasia of Animals (2020). Following assessment of adequate depth of anesthesia by eye-blink reflexes and toe-pinch withdrawal, the thorax was opened and the right atrium was cut to allow drainage. The heart was perfused with 20 mL of ice-cold phosphate-buffered saline (PBS), followed by 20 mL of ice-cold 4% paraformaldehyde (1.04005.1000, Merck Millipore) in 0.1 M phosphate buffer (PB; pH 7.4). Brains were removed from the skull, postfixed in the same fixative overnight and cryoprotected in 30% sucrose in 0.1 M PB at 4 °C. Fixed brains of male *App*^*NL-F/NL-F*^ (17) mice (RBRC06343, RIKEN BioResource Research Center) at 14 months old were provided by Dr. A. Miyawaki (RIKEN Center for Brain Science). The brains were mounted on a freezing microtome (REM-710; Yamato Kohki Industrial) and cut into 30-µm-thick coronal sections (18).

### IHC staining

IHC was performed by a free-floating method at 20–25 °C (19). The primary and secondary Abs used are listed in S1 and S2 Tables respectively. Abs and streptavidin were diluted in 0.3% Triton X-100 in PBS (PBS-X) containing 0.12% λ-carrageenan (035-09693; Sigma-Aldrich) and 1% normal donkey serum (S30-100ML, Merck Millipore). Heat induced epitope retrieval (HIER) was accomplished by overnight incubation at 60 °C in a citrate buffer (ab93678, abcam) prior to IHC staining. Brain sections immunostained for Aβ were mounted onto glass slides (Superfrost micro slide glass APS-coated, Matsunami Glass) and coverslipped with Fluoromount/Plus™ (K048, Diagnostic Biosystems).

For single IHC staining with anti-Aβ Abs, brain sections were washed twice for 10 min in PBS-X and reacted overnight with primary Abs. The sections were washed twice for 10 min in PBS-X and reacted for 2 h with secondary Abs. After washing twice for 10 min in PBS-X, the sections were incubated with PBS-X containing FSB (10 or 20 µg/mL, F308, Dojindo) and NeuroTrace 530/615 Red Fluorescent Nissl Stain (1:100, N21482, Thermo Fisher Scientific), and washed twice for 10 min in PBS-X.

For double IHC staining with 6E10 and 82E1 or D54D2 Aβ Abs, brain sections were reacted first with 6E10 Ab, and then with biotinylated 82E1 or D54D2 Abs. In double IHC with 6E10 and 82E1 mouse monoclonal Abs, Alexa Fluor (AF) 488-conjugated Fab fragments of goat ant-mouse IgG1 Ab (8.0 µg/mL) was applied prior to incubation with the biotinylated 82E1 Ab. AF 568-conjugated streptavidin (2 µg/mL, S11226, Thermo Fisher Scientific) was used for the detection of the biotinylated anti-Aβ Ab. In double IHC with 6E10 and D54D2 Aβ Abs, secondary Abs were simultaneously applied following incubation with D54D2 Ab. After IHC staining, brain sections were stained with FSB as above.

Fluorochromized Tyramide-Glucose Oxidase (FT-GO) immunofluorescence (IF) was adapted for sensitive detection of 6E10 Aβ Ab (19). Following incubation for 30 min in PBS containing 1% hydrogen peroxide, brain sections were washed twice for 10 min in PBS-X and reacted overnight with biotinylated 6E10 Ab. The sections were washed twice for 10 min PBS-X, incubated for 2 h in avidin-biotin complex solution (1:500, PK-6100, Vectastain Elite ABC kit, Vector Laboratories), and washed twice for 10 min in PBS-X and twice for 5 min in 0.1 M PB. Then, the sections were incubated for 10 min with an FT-GO reaction mixture which contained 10 μM CF568 tyramide (92173, Biotium). FT-GO reaction was initiated by adding β-D-glucose (16804-32, Nacalai Tesque) into the mixture at 2 mg/mL and proceeded for 30 min. Following two washes in PBS-X for 10 min each, the sections were reacted overnight with a complex of 82E1 Ab (1.0 µg/mL) and AF 488-conjugated Fab fragments of goat ant-mouse IgG1 Ab (1.0 µg/mL) and washed twice for 10 min in PBS-X. The primary and secondary Ab complex was prepared following the manufacturer instruction. In the indirect detection compared with FT-GO signal amplification, unconjugated 6E10 Ab and Rhodamine Red X-conjugated Fab fragments of goat ant-mouse IgG1 Ab were used as primary and secondary Abs, respectively. Brain sections were reacted with these Abs prior to incubation with 82E1 Ab.

### Image acquisition and processing

Images were captured with a confocal laser scanning microscope (TCS SP8; Leica Microsystem). A 16× multi-immersion (HC FLUOTAR 16x/0.60 IMM CORR VISIR, NA = 0.60, Leica Microsystems) and 25× water-immersion (HC FLUOTAR L 25x/0.95 W VISIR, NA = 0.95, Leica Microsystems) objective lenses were used. The confocal pinhole was adjusted to 1.0 or 2.0 Airy unit. Z-stack images were collected at 1.5 to 8.0 µm intervals at 512×512 or 1,024×1,024 pixel resolutions. Captured images were stitched and processed to create maximum intensity projection images using Leica Application Suite X software (LAS X, ver. 3.5.5.19976, Leica Microsystems). The brightness and contrast of images were adjusted using Fiji software (20) (ver. 2.14.0/1.54 f).

### Quantification

To assess IHC reactivity of 6E10, 82E1 and D54D2 Aβ Abs, immunohistochemically stained area (IHCSA) of each Ab was measured in the cerebral cortex and hippocampus using Fiji software. After subtracting background noise, the number of pixels above a threshold was counted and designated as the IHCSA. The threshold was set manually for each image. The IHCSA was normalized to the FSB-stained area, a proxy for Aβ load in each brain section. In the cerebral cortex, the IHCSA was analyzed in the region which is located medial to the outer edge of CA3 pyramidal cell layer and lateral to the inner edge of the dentate gyrus (DG) granule cell layer. The hippocampus was analyzed in its entirety. For direct comparisons of staining properties between 6E10 and 82E1 or D54D2 Aβ Abs, FSB-labeled plaques were randomly selected from the cerebral cortex of *App*^*NL-F/NL-F*^ mice.

To assess effects of signal amplification resulting from FT-GO IF and antigen unmasking with HIER, IHCSA of 6E10 or 82E1 Aβ Abs was measured and quantified in the cerebral cortex of *App*^*NL-F/NL-F*^ mice as above. The coverage area of 82E1 IHCSA by 6E10 IHCSA was calculated using Fiji software. The region medial to the outer edge of CA3 pyramidal cell layer and lateral to the inner edge of the DG granule cell layer was subjected to analysis.

### Statistical analysis

Statistical analyses were conducted using GraphPad Prism 9 software (Version 9.4.1 (458), GraphPad Software). For comparisons among independent groups, Brown-Forsythe and Welch one-way analysis of variance (ANOVA) followed by Tukey-Kramer post hoc test (*App*^*NL-G-F/NL-G-F*^ mouse hippocampus) or Kruskal-Wallis test followed by Dunn’s post hoc test (*App*^*NL-F/NL-F*^ mouse cerebral cortex and hippocampus, and *App*^*NL-G-F/NL-G-F*^ mouse cerebral cortex) were used. Wilcoxon test was used for paired comparisons between groups (Direct comparisons between 6E10 and 82E1 or D54D2 Abs). For comparisons between two independent groups with a normal distribution, unpaired t test was used (FT-GO signal amplification and antigen unmasking using HIER). All tests were two-sided. Statistical significance was set at *P* < 0.05. Graphed data are represented as means ± standard deviations (SDs).

IHCSA of 6E10, 82E1 and D54D2 Aβ Abs were examined in seven to nine sections from three *App*^*NL-F/NL-F*^ and *App*^*NL-G-F/NL-G-F*^ mice. Thirty plaques in nine brain sections from three *App*^*NL-F/NL-F*^ mice were used for direct comparisons between 6E10 and 82E1 or D54D2 Aβ Abs. Nine sections from three *App*^*NL-F/NL-F*^ mice were used to evaluate signal enhancement in 6E10 FT-GO IF and antigen unmasking using HIER. The exact values of n are stated in the corresponding figure legends. Unusual outliers were not found in the datasets.

### Data availability

The datasets generated during and/or analyzed during the current study and all biological materials reported in this article are available from the corresponding authors on reasonable request.

## Results

### Differential IHC reactivity of N-terminal Aβ Abs

We examined IHC reactivity of monoclonal Abs (mAbs) against N-terminal regions of human Aβ peptide, because 1) N-terminal Aβ Abs are assumed to detect total Aβ peptide in the specimen, 2) diversity of Aβ species generated by γ-secretase cleavage affects IHC outcomes using anti-Aβ Abs against the C-terminal residues (13) and 3) FAD mutations located at the middle of the Aβ sequence can alter binding properties of anti-Aβ Abs that react the region(17). The Aβ Abs studied were 6E10 (21), 82E1 (22) and D54D2 Aβ monoclonal Abs, which all were generated by immunization with N-terminal residues of Aβ peptide and commercially available. Table 1 summarizes host animals, isotypes, immunogens, epitopes and suppliers of these anti-Aβ Abs.

**Table 1.** Aβ Ab information used in this study.

| Clone name | Host/Isotype | Immunogen | Epitope | Manufacturer, Cat. No. | RRID |
| --- | --- | --- | --- | --- | --- |
| 6E10(21) | Mouse/IgG1 | Human Aβ 1-16 peptide | Aβ 3–8 peptide | BioLegend, 80301 | AB_2565328 |
| 82E1(22) | Mouse/IgG1 | Human Aβ 1-16 peptide | N-terminus end<br>(Aβ 1–5 peptide)(22) | Immuno-Biological<br>Laboratories, 10323 | AB_1630806 |
| D54D2 | Rabbit/IgG | Near N-terminus of<br>human Aβ peptide | Not determined | Cell Signaling<br>Technology, 8243S | AB_2797642 |

We first compared IHC reactivity of these three Abs in *App*^*NL-G-F*^ mouse model (17) (Fig. 1a). Aβ IHC using 6E10, 82E1 and D54D2 Aβ Abs was carried out in frozen brain sections of *App*^*NL-G-F/NL-G-F*^ mice. These brain sections were further stained with a Congo-red derivative, 1-Fluoro-2,5-bis(3-carboxy-4-hydroxystyryl)benzene (FSB) (23), to assess Aβ deposition in each brain section. IHC reactivity for Aβ peptides was quite different among 6E10, 82E1 and D54D2 Abs (Fig. 1a). While moderate and scattered immunoreactivity was found in brain sections stained with 6E10 Ab (Fig. 1a_1_), 82E1 and D54D2 Abs labeled more massively and intensely brain sections of *App*^*NL-G-F/NL-G-F*^ mice (Fig. 1a_2, 3_). IHC reactivity were similar between 82E1 and D54D2 Aβ Abs. The same tendency was found in *App*^*NL-F/NL-F*^ mouse models (Fig. 1b), where Aβ burden is much less severe than *App*^*NL-G-F/NL-G-F*^ mice (17). IHCSA normalized to Aβ fibril deposition assessed by FSB labeling was measured and quantified in the cerebral cortex and hippocampus of both models (Fig. 1c, d). IHCSA of 82E1 and D54D2 Aβ Abs was 2.49 to 4.49 times broader than that of 6E10 Aβ Ab. There were no significant differences in IHCSA between 82E1 and D54D2 Abs (Fig. 1c, d). Additionally, the significant difference in IHCSA was not detected between 6E10 and 82E1 Abs in the hippocampus of *App*^*NL-F/NL-F*^ mice.

**Fig 1.**
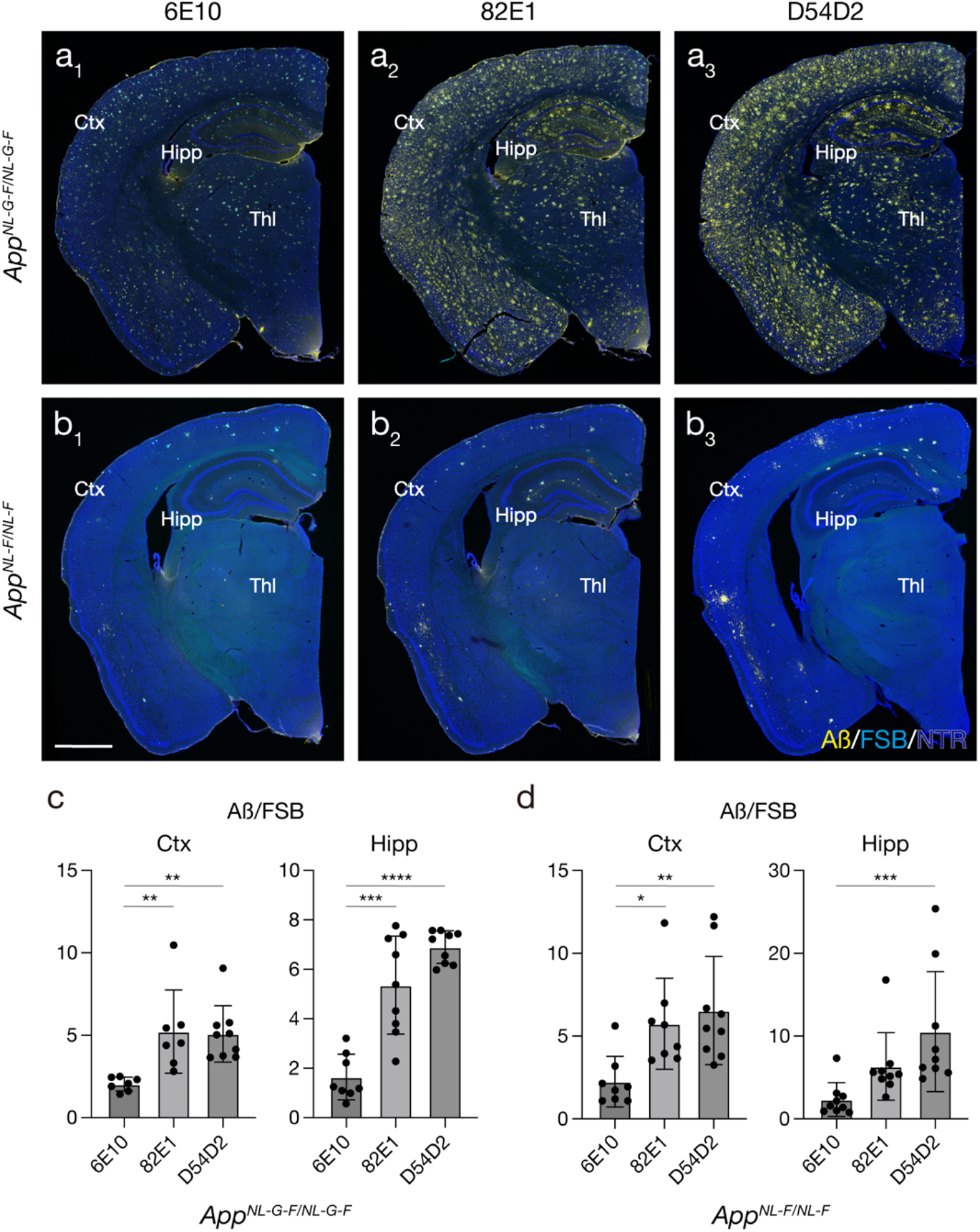
Differential IHC reactivity among 6E10, 82E1 and D54D2 Aβ Abs. **a, b**) Aβ IHC in *App*^*NL-G-F/NL-G-F*^ (**a**) and *App*^*NL-F/NL-F*^ (**b**) mouse brain sections. 6E10 (**a**_**1**_, **b**_**1**_), 82E1 (**a**_**2**_, **b**_**2**_) and D54D2 (**a**_**3**_, **b**_**3**_) Aβ Abs are used for the IHC. **c, d**) Histograms showing The IHCSA of 6E10, 82E1 and D54D2 Abs in *App*^*NL-G-F/NL-G-F*^ (**c**) and *App*^*NL-F/NL-F*^ (**d**) mice. The IHCSA in the cerebral cortex (left) and hippocampus (right) are represented (*App*^*NL-G-F/NL-G-F*^: n = 7 sections, 6E10 Ab in the Ctx; n = 7 sections, 82E1 Ab in the Ctx; n = 9 sections, D54D2 Ab in the Ctx from 3 animals; *H* = 14.59, *df* = 2, *P* = 0.0009, Kruskal-Wallis test; n = 8 sections, 6E10 Ab in the Hipp; n = 9 sections, 82E1 Ab in the Hipp; n = 9 sections, D54D2 Ab in the Hipp from 3 animals; *F* = 35.42, *df* = 2, *P* < 0.0001, Brown-Forsythe ANOVA test. *App*^*NL-F/NL-F*^: n = 8 sections, 6E10 Ab in the Ctx; n = 8 sections, 82E1 Ab in the Ctx; n = 9 sections, D54D2 Ab in the Ctx from 3 animals; *H* = 11.88, *df* = 2, *P* = 0.0026, Kruskal-Wallis test; n = 9 sections, 6E10 Ab in the Hipp; n = 9 sections, 82E1 Ab in the Hipp; n = 9 sections, D54D2 Ab in the Hipp from 3 animals; *H* = 13.91, *df* = 2, *P* = 0.001, Kruskal-Wallis test). Data are represented as means ± SDs. Ctx: cerebral cortex, Hipp: hippocampus, Thl: Thalamus. Scale bar: 1000 µm. \**P* < 0.05; \*\**P* < 0.01; \*\*\**P* < 0.001; \*\*\*\**P* < 0.0001.

We then directly compared IHC reactivity of the anti-Aβ Abs by double IHC staining (Fig. 2). We analyzed IHC reactivity in the cerebral cortex of *App*^*NL-F/NL-F*^ mice, because massive Aβ plaque burden in *App*^*NL-G-F/NL-G-F*^ mice makes it difficult to examine staining properties of these Abs in detail (Fig. 1a). Given that 82E1 and D54D2 Abs showed comparable IHC reactivity (Fig. 1c, d), IHC reactivity was compared between 6E10 and 82E1 or D54D2 N-terminal Aβ Abs. For this, we conducted sequential double IHC staining. Following incubation of 6E10 Ab, 82E1 or D54D2 Ab was applied to brain sections to prevent the latter two Abs from preoccupying the epitope of 6E10 Ab. Again, 82E1 and D54D2 Abs showed much stronger IHC reactivity than 6E10 Ab in the direct comparison (Fig. 2a, b). Of all analyzed plaques, 82E1 and D54D2 Abs stained broader regions than 6E10 Ab did (Fig. 2c, d). IHCSA of 82E1 and D54D2 Abs was 2.67 and 4.25 times broader respectively than that of 6E10 Ab (Fig. 2c, d). Whereas 6E10 immunoreactivity was mainly localized to plaque cores, 82E1 and D54D2 Abs stained much broader regions over amyloid cores visualized by FSB labeling (Fig. 2a, b). Interestingly, 82E1 and D54D2 Abs showed blurred filamentous immunoreactivity beyond plaque cores (Fig. 2a, b). 6E10 IHCSA narrower than the FSB-stained area was found in approximately half of analyzed plaques (33 plaques in 60 plaques).

**Fig 2.**
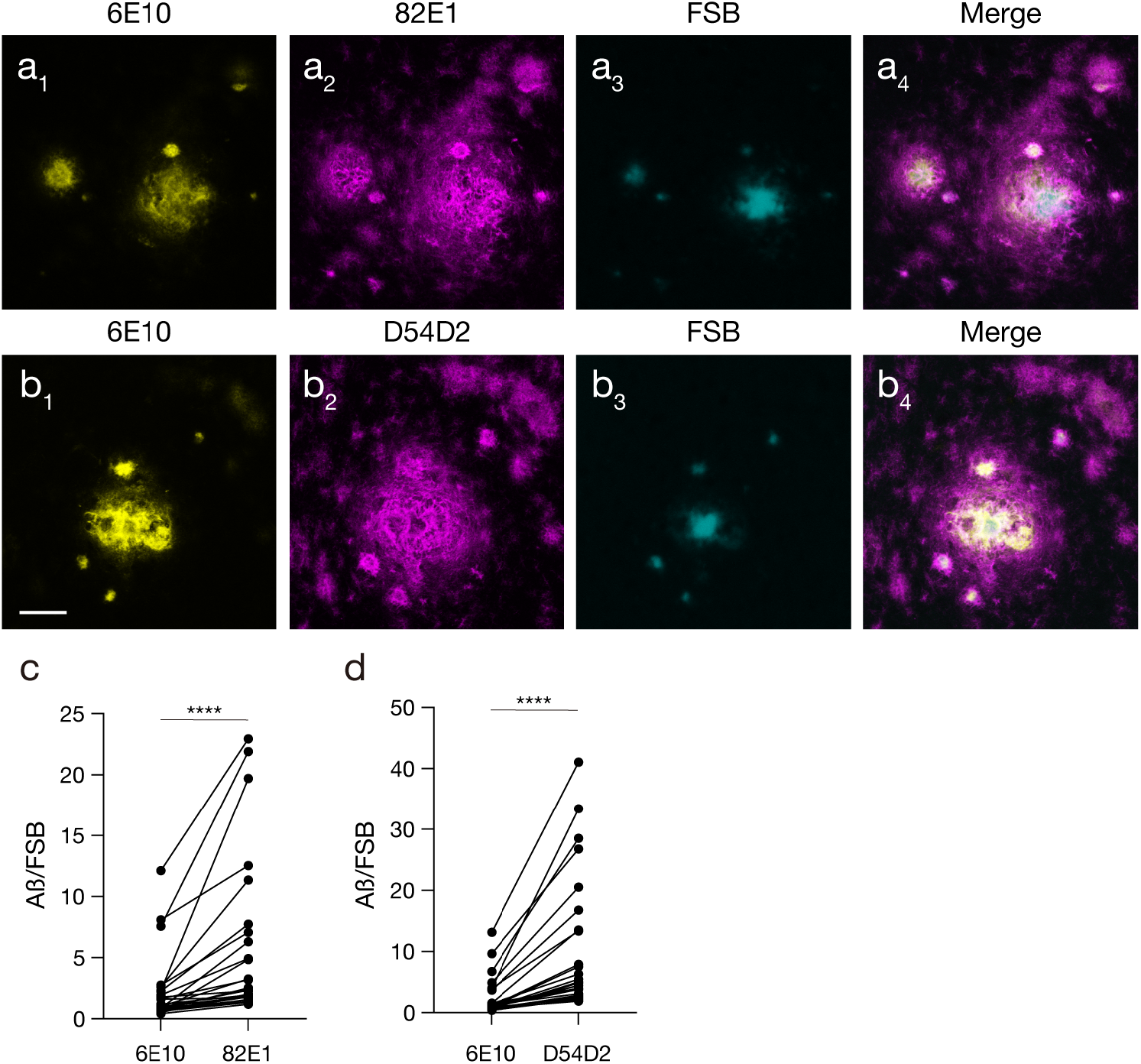
Direct comparisons of IHC characteristics between 6E10 and 82E1 or D54D2 Aβ Abs. **a, b**) Double IHC with 6E10 and 82E1 (**a**) or D54D2 (**b**) Aβ Abs in the cerebral cortex of *App*^*NL-F/NL-F*^ mice. FSB is used for labeling Aβ fibrils. **a**_**1-4**,_ **b**_**1-4**_) Representative images for 6E10 Ab (yellow, **a**_**1**_, **b**_**2**_), 82E1 (magenta, **a**_**2**_) or D54D2 Ab (magenta, **b**_**2**_) and FSB (cyan, **a**_**3**_, **b**_**3**_) staining. **a**_**4**_, **b**_**4**_) Merged images of (**a**_**1**_), (**a**_**2**_) and (**a**_**3**_), and (**b**_**1**_), (**b**_**2**_) and (**b**_**3**_), respectively. **c, d**) Direct comparisons of IHCSA of 6E10 and 82E1 (**c**) or D54D2 Aβ Abs (**d**) (n = 30 plaques from 9 sections of 3 animals for each; 6E10 vs 82E1 Abs: *W* = 465.0, *df* = 1, *P* < 0.0001, Wilcoxon matched-pairs signed rank test; 6E10 vs D54D2 Abs: *W* = 465.0, *df* = 1, *P* < 0.0001, Wilcoxon matched-pairs signed rank test). Data are represented as means ± SDs. Scale bar: 30 µm. \*\*\*\**P* < 0.0001.

One possible explanation for the differential IHC outcomes among 6E10, 82E1 and D54D2 N-terminal Aβ Abs is the difference of IHC sensitivity of these Abs. If differential IHC sensitivity underlies the differential outcomes of IHC, signal amplification techniques that enhances IHC sensitivity should close the gap of IHC reactivity among Abs. We thus conducted FT-GO IF for 6E10 Ab and compared IHC reactivity of 82E1 Ab. FT-GO is a fluorescent tyramide signal amplification system (TSA) technique, yielding 10- to 30-fold signal amplification compared with indirect IHC detection (19). Because the epitope of D54D2 Ab has not been described in detail (Table 1), this Ab was excluded from the analysis. Even after signal amplification by FT-GO, IHCSA of 82E1 Ab was still much broader than that of 6E10 Ab (6E10/82E1^+^ IHCSA in 82E1^+^ IHCSA [means ± SDs]: FT-GO, 0.442 ± 0.081) (Fig. 3). FT-GO signal amplification actually enhanced IHC sensitivity of 6E10 Ab: IHCSA of FT-GO detection was significantly broader than that of indirect detection (Fig. 3c) (6E10/82E1^+^ IHCSA in 82E1^+^ IHCSA [means ± SDs]: indirect method, 0.318 ± 0.079; FT-GO, 0.442 ± 0.081, *P* = 0.0046, unpaired t test). These results imply that the difference of IHC sensitivity is only partially attributable to the differential IHC outcomes between 6E10 and 82E1 Abs. We further examined antigen masking effects on the differential IHC reactivity. Antigen retrieval techniques such as HIER is frequently used for Aβ IHC staining to unmask epitopes. However, an HIER procedure did not affect differential IHC outcomes between 6E10 and 82E1 Abs (Fig. 4). The gap of IHC reactivity between 6E10 and 82E1 Abs still remained in brain sections treated with HIER (Fig. 4c) (6E10/82E1^+^ IHCSA in 82E1^+^ IHCSA [means ± SDs]: wo/HIER, 0.362 ± 0.087; w/HIER, 0.407 ± 0.073, *P* = 0.2454, unpaired t test).

**Fig 3.**
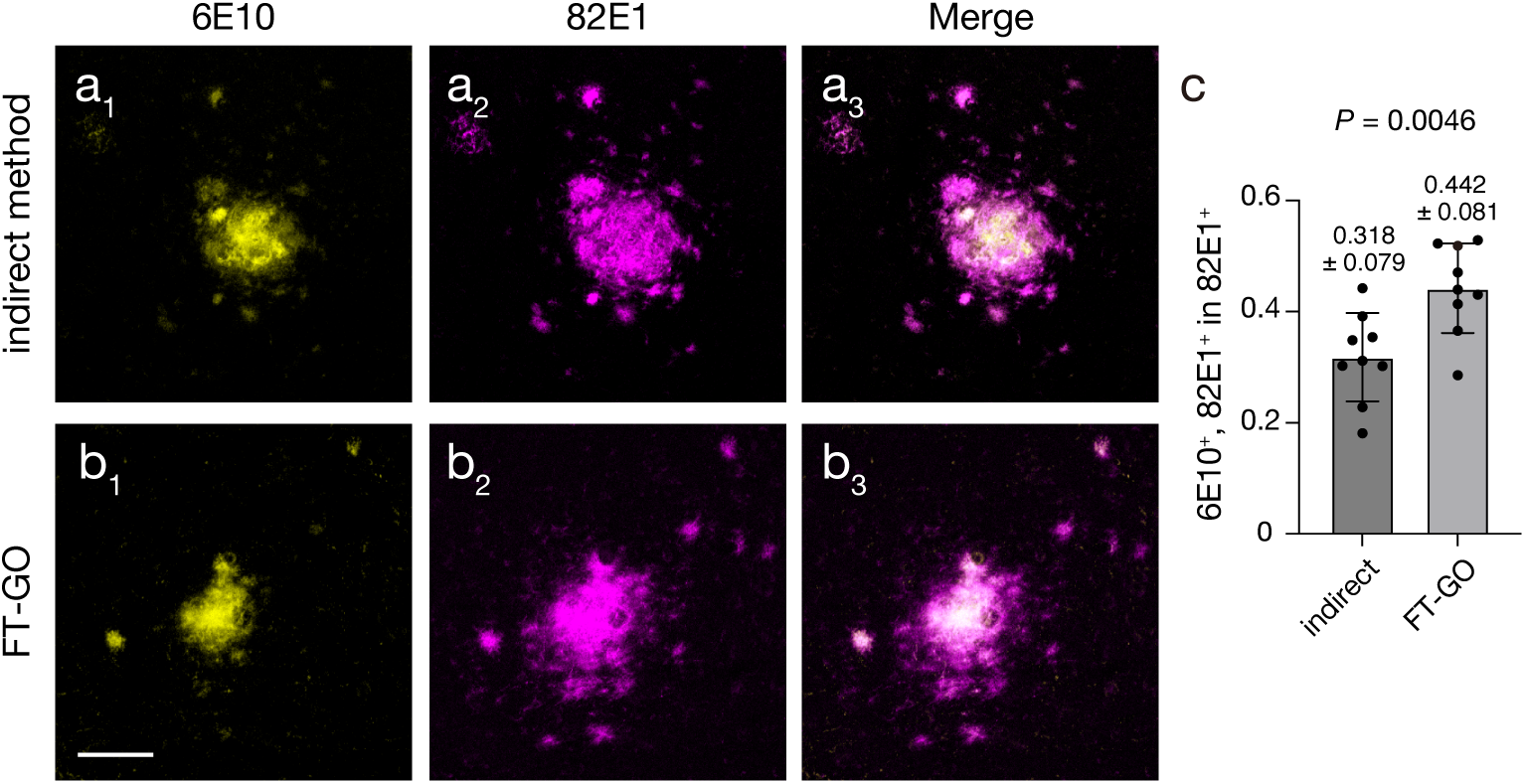
Signal amplification characteristics of 6E10 Aβ Ab by FT-GO. **a, b**) Double IHC with 6E10 and 82E1 Aβ Abs in the cerebral cortex of *App*^*NL-F/NL-F*^ mice. 6E10 (**a**) and Biotin-6E10 Ab (**b**) are detected with an indirect method (**a**) and FT-GO signal amplification (**b**), respectively. 82E1 Ab is labeled with an AF 488 conjugated FabuLight antibody prior to incubation with brain sections. **a**_**1-3**,_ **b**_**1-3**_) Representative images for 6E10 Ab (yellow, **a**_**1**_, **b**_**1**_) and 82E1 Ab (magenta, **a**_**2**_, **b**_**2**_) staining. **a**_**3**_, **b**_**3**_) Merged images of (**a**_**1**_) and (**a**_**2**_), and (**b**_**1**_) and (**b**_**2**_), respectively. **c**) Histograms representing the proportion of 82E1 IHCSA covered by 6E10 IHCSA (n = 9 sections from 3 animals for each; *t* = 3.288, *df* = 16, *P* = 0.0046, unpaired t test). Data are represented as means ± SDs. Scale bar: 50 µm.

**Fig 4.**
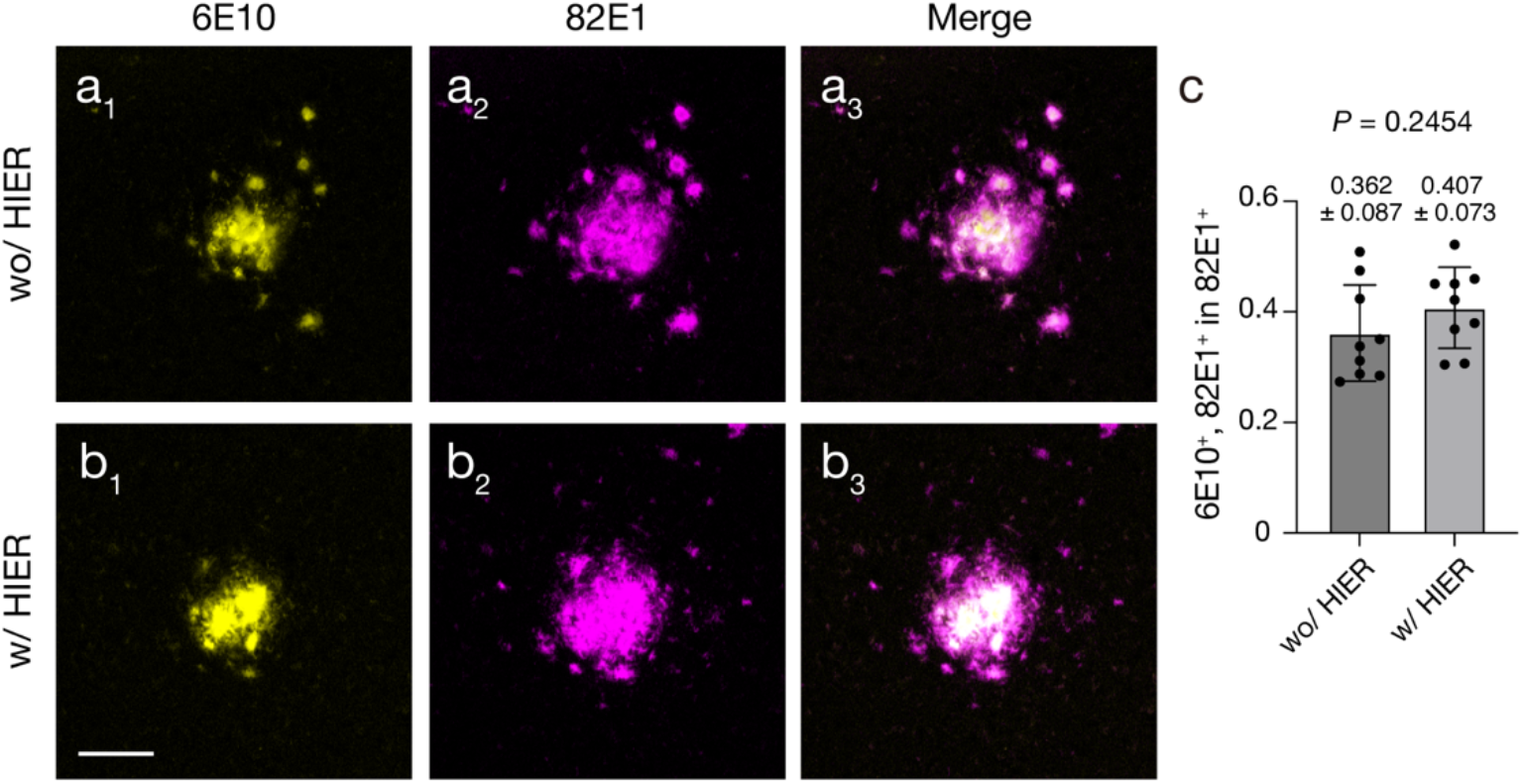
The effect of antigen unmasking on the differential IHC reactivity between 6E10 and 82E1 Aβ Abs. **a, b**) Double IHC with 6E10 and 82E1 Aβ Abs in the cerebral cortex of *App*^*NL-F/NL-F*^ mice. Double Aβ IHC is performed in non-treated (**a**) and HIER-treated (**b**) brain sections. **a**_**1-3**,_ **b**_**1-3**_) Representative images for 6E10 Ab (yellow, **a**_**1**_, **b**_**1**_) and 82E1 Ab (magenta, **a**_**2**_, **b**_**2**_) staining. **a**_**3**_, **b**_**3**_) Merged images of (**a**_**1**_) and (**a**_**2**_), and (**b**_**1**_) and (**b**_**2**_), respectively. **c**) Histograms representing the proportion of 82E1 IHCSA covered by 6E10 IHCSA (n = 9 sections from 3 animals for each; *t* = 1.206, *df* = 16, *P* = 0.2454, unpaired t test). Data are represented as means ± SDs. Scale bar: 50 µm.

## Discussion

In the present study, we analyzed IHC characteristics of two mouse and one rabbit monoclonal Abs against N-terminal regions of Aβ peptide that recognize several isoforms of Aβ peptide and transgenically expressed human APP. We found significant differences of IHC reactivity among these Abs in both *App*^*NL-F*^ and *App*^*NL-G-F*^ mouse models: while 6E10 immunosignal was mainly localized to plaque cores labeled by FSB, 82E1 and D54D2 Abs stained much more broadly beyond cores. Our results further implied that the differential IHC outcomes between 6E10 and 82E1 Abs was only partially attributable to IHC sensitivity of these Abs by using a fluorescent TSA system, FT-GO. Antigen unmasking with a HIER technique did not affect the difference of IHC reactivity between 6E10 and 82E1 Abs.

6E10, 82E1 and D54D2 N-terminal Aβ Abs showed significant differences regarding their IHC characteristics, even despite similarities in their prepared immunogens. These Aβ Abs are widely assumed and used to detect total Aβ peptides. IHCSA of 82E1 and D54D2 Abs was much broader than that of 6E10 Ab (Fig. 1). In contrast to immunoreactivity of 6E10 Ab, which was mainly localized to amyloid cores, 82E1 and D54D2 Abs showed blurred filamentous immunosignals beyond plaque cores in addition to immunoreactivity against core region (Fig. 2). The IHC reactivity against filamentous materials around plaque cores can account for the broader IHCSA of 82E1 and D54D2 Abs. The blurred filamentous immunoreactivity might reflect the deposition of protofibrillar and/or fibrillar forms of Aβ peptides onto the brain (2, 3). Our finding that the coverage area of 82E1 IHCSA by 6E10 immunosignals was increased by FT-GO signal amplification indicates that differential IHC sensitivity between 6E10 and 82E1 Abs partially explains their differential IHC outcomes (Fig. 3c). However, the differential IHC sensitivity does not account for most of the difference of IHC reactivity between these two Aβ Abs, because a substantial portion of 82E1 IHCSA was still not covered by 6E10 immunosignal even after FT-GO signal amplification (Fig. 3b). Our finding that an HIER procedure failed to fill the gap of IHC reactivity between 6E10 and 82E1 Abs argues against the view that antigen masking effects of formaldehyde fixation causes the differential IHC reactivity between the two Aβ Abs (Fig. 4). The slight difference of epitopes within N-terminal regions of Aβ peptide between 6E10 and 82E1 Aβ Abs (Table 1) might generate and/or lose affinities for specific forms of the peptides in frozen brain sections. While the epitope of 6E10 Ab lies within amino acids 3-8 of Aβ peptide, 82E1 Ab is an N-terminus end specific antibody (22). We cannot rule out the possibility that stronger signal amplification than FT-GO close the gap completely between 6E10 and 82E1 IHCSA.

We found differential IHC characteristics of anti-Aβ Abs in both *App*^*NL-G--F*^ and *App*^*NL-F*^ mouse models (Fig. 1). *App*^*NL-G-F*^ model which harbors Swedish (KM670/671NL), Beyreuther/Iberian (I716F) and Arctic (E693G) mutations is being used most frequently as it develops Aβ pathology much faster than other *App* knock-in lines (24). Aβ sequence of *App*^*NL-F*^ model, which carries the Swedish and Beyreuther/Iberian mutations, is identical to that of wild-type human (17). We consider that our findings of significant differences of IHC characteristics among 6E10, 82E1 and D54D2 Abs could be extrapolatable to other mouse models of AD which express APP FAD mutants transgenically, because the most commonly used mutations in these FAD mutant models are the Swedish and/or Beyreuther/Iberian mutations that are carried by *App*^*NL-G-F*^ and *App*^*NL-F*^ mouse models (25). Notably, Aβ42 filament structure in *App*^*NL-F*^ model are identical to type II filaments from human brains (26). Further studies are required to determine whether the difference of IHC reactivity among Aβ Abs is also found in the brain of AD patients.

## Conclusions

A wide variety of anti-Aβ Abs developed over the past decades make the knowledge about IHC reactivity of Abs against Aβ peptides important for accurate and reliable evaluation of Aβ plaque burden in patients as well as models of AD. Our results demonstrate that Aβ IHC outcomes highly depend on what kind of Ab is used for the analysis. This could be of importance for meta-analysis where the results of Aβ IHC staining using various kinds of Abs are pooled.

## Acknowledgements

We thank Drs. Hiroshi Hama (RIKEN Center for Brain Science) and Atsushi Miyawaki (RIKEN Center for Brain Science) for their kind provision of *App*^*NL-F/NL-F*^ mouse brains; and Drs. Takashi Saito (Nagoya City University) and Takaomi C Saido (RIKEN Center for Brain Science) for sharing *App*^*NL-G-F*^ mice. K.A. is supported by Juntendo Specialized Training for Aspiring Researchers (J-STAR) program at Juntendo University School of Medicine.

## Author contributions

Conceptualization, K.A., K.Y. and S.I.; Methodology, K.A., K.Y. and S.I.; Investigation, K.A., K.Y., S.I. and H.H.; Formal Analysis, K.A., K.Y. and S.I.; Writing – Original Draft, K.Y.; Writing – Review & Editing, K.A., K.Y., S.I., M.K. and H.H.; Visualization, K.A., and K.Y.; Project Administration, K.Y.; Funding Acquisition, K.Y., S.I., M.K. and H.H..

## Supporting information

**S1 table. Primary Abs used in this study**.

**S2 table. Secondary Abs used in this study**

## Notes

### Competing Interest Statement

The authors have declared no competing interest.

### Summary of Updates

The supplemental files have been uploaded. No changes have been made to the main text.

