## Supplementary material for "The difference in immunohistochemical reactivity of monoclonal antibodies against amino-terminal residues of amyloid-β peptide": S1 Table

### S1 Table: Primary Abs used in this study

| Antibody name | Host/Isotype | Clone | Manufacturer, Cat. No. | RRID | Concentration |
|---|---|---|---|---|---|
| Anti-β-Amyloid, 1-16 Antibody | Mouse/IgG1 | 6E10 | BioLegend, 80301 | AB_2565328 | 1.0 μg/ml |
| Biotin anti-β-Amyloid, 1-16 Antibody | Mouse/IgG1 | 6E10 | BioLegend, 80307 | AB_2564656 | 2.0 μg/ml |
| Anti-Human Amyloidβ (N) (82E1) Mouse IgG MoAb | Mouse/IgG1 | 82E1 | Immuno-Biological Laboratories, 10323 | AB_1630806 | 1.0 μg/ml |
| Anti-Human Amyloidβ (N) (82E1) Mouse IgG MoAb Biotin | Mouse/IgG1 | 82E1 | Immuno-Biological Laboratories, 10326 | AB_2341281 | 1.0 μg/ml |
| β-Amyloid (D54D2) XP® Rabbit mAb | Rabbit/IgG | D54D2 | Cell Signaling Technology, 8243S | AB_2797642 | 1:800 |
