## Supplementary material for "The difference in immunohistochemical reactivity of monoclonal antibodies against amino-terminal residues of amyloid-β peptide": S2 Table

### S2 Table: Secondary Abs used in this study

| Antibody name | Antibody format | Manufacturer, Cat. No. | RRID | Concentration |
|---|---|---|---|---|
| Donkey anti-Mouse IgG (H+L) Highly Cross-Adsorbed Secondary Antibody, Alexa Fluor™ 488 | Whole IgG | Thermo Fisher Scientific, A-21202 | AB_141607 | 10 μg/ml |
| Alexa Fluor® 488 AffiniPure Fab Fragment Goat Anti-Mouse IgG1, Fcγ fragment specific | Fab fragment | Jackson ImmunoResearch, 115-547-185 | AB_2632534 | 1.0 or 8.0 μg/ml |
| Donkey anti-Rabbit IgG (H+L) Highly Cross-Adsorbed Secondary Antibody, Alexa Fluor™ 488 | Whole IgG | Thermo Fisher Scientific, A-21206 | AB_2535792 | 10 μg/ml |
| Donkey anti-Rabbit IgG (H+L) Highly Cross-Adsorbed Secondary Antibody, Alexa Fluor™ 568 | Whole IgG | Thermo Fisher Scientific, A10042 | AB_2534017 | 10 μg/ml |
| Rhodamine Red-X AffiniPure Fab Fragment Goat Anti-Mouse IgG1, Fcγ fragment specific | Fab fragment | Jackson ImmunoResearch, 115-297-185 | AB_2632519 | 28 μg/ml |
